# From the Andes to the desert: First overview of the bacterial community in the Rimac river, the main source of water for Lima, Peru

**DOI:** 10.1101/2020.08.16.252965

**Authors:** Pedro E. Romero, Erika Calla-Quispe, Camila Castillo-Vilcahuaman, Mateo Yokoo, Hammerly Lino Fuentes-Rivera, Jorge L. Ramirez, Alfredo J. Ibáñez, Paolo Wong

## Abstract

**Background:** The Rimac river is the main source of water for Lima, Peru’s capital megacity. The river is constantly affected by different types of contamination including mine tailings in the Andes and urban sewage in the metropolitan area. We aim to produce the first characterization of bacterial communities in the Rimac river using a 16S rRNA amplicon sequencing approach which would be useful to identify bacterial diversity and potential understudied pathogens.

**Results:** We report a higher diversity in bacterial communities from the Upper and, especially, Middle Rimac compared to the Lower Rimac (Metropolitan zone). Samples were generally grouped according to their geographical location. Bacterial classes Alphaproteobacteria, Bacteroidia, Campylobacteria, Fusobacteriia, and Gammaproteobacteria were the most frequent along the river. *Arcobacter cryaerophilus* (Campylobacteria) was the most frequent species in the Lower Rimac while *Flavobacterium succinicans* (Bacteroidia) and *Hypnocyclicus* (Fusobacteriia) were the most predominant in the Upper Rimac. Predicted metabolic functions in the microbiota include bacterial motility, quorum sensing and xenobiotics metabolism. Additional metabolomic analyses showed the presence natural flavonoids and antibiotics in the Upper Rimac, and herbicides in the Lower Rimac.

**Conclusions:** The dominance in the Metropolitan area of *Arcobacter cryaerophilus*, an emergent pathogen associated with fecal contamination and antibiotic multiresistance, but that is not usually reported in traditional microbiological quality assessments, highlights the necessity to apply next-generation sequencing tools to improve pathogen surveillance. We believe that our study will encourage the integration of omics sciences in Peru and its application on current environmental and public health issues.

## 1. Introduction

Worldwide, water quality problems are associated with poverty conditions and lack of efficient sanitation, especially in developing countries [1]. These problems are recurrent in highly populated cities where most of the waste is directly washed to nearby rivers [2]. Lima, the capital city of Peru, is the second largest desert city in the world [3]. Its Metropolitan area is inhabited by more than 10 million people creating enormous challenges for environmental and public health [4]. For instance, it has been shown that water quality in Lima is a significant risk factor for pathogenic infections in children [5].

The Rimac river is the main source of drinking water for the Lima Metropolitan area. Recently, more than 700 pollution sources were identified by the National Authority of Water [6]. The river is constantly polluted by mine tailings in its Upper zone close to the central Peruvian Andes, by agricultural wastewater in its Middle zone, and by industrial wastewater and urban sewage in its Lower zone within the Metropolitan area nearby the Pacific Ocean [7]. The lack of an efficient wastewater treatment in the Metropolitan region promotes the presence of potentially pathogenic bacteria such as *Escherichia coli, Salmonella typhi*, or *Vibrio cholerae*, associated with fever and diarrhea symptoms [2,8].

Traditionally, assessments of water quality and bacterial contamination in Peru are focused on evaluating the presence of common coliforms (e.g. *Citrobacter, Escherichia, Enterobacter, Klebsiella*) [9]. However, there are a plethora of likely pathogens that are not currently studied using classic methods because of taxonomic assignation problems or the existence of non-culturable phenotypes [10]. In recent years, advances in the study of bacterial communities and diversity have occurred because of the improvement of next-generation sequencing (NGS) methodologies. One of these techniques, 16S rRNA amplicon sequencing, uses a fragment of the 16S ribosomal gene to obtain a diversity profile of the bacterial community in a specific environment. Therefore, it has been used to study different types of samples from the internal microbiota of several species, including humans to environmental community surveys [11].

16S rRNA amplicon sequencing has been also used to study bacterial communities from several rivers. For instance, a study in the Danube river, which crosses many countries from central Europe, found a higher microbial community richness in the Upper basin, while in the Lower basin, there was a predominance of only a few free-living and particle associated bacteria [12]. Studies in rivers have also looked for the occurrence of potential pathogens. A report from the river Tama (Tokyo, Japan) showed that the predominant bacteria genus was *Flavobacterium* (Bacteroidia), a freshwater fish pathogen [13]. Recently, a study in the Pinheiros river (Sao Paulo, Brazil), one of the most polluted Brazilian rivers, found the predominance of *Arcobacter cryaerophylus* (Campylobacteria). This species is considered an emerging pathogen and an indicator of fecal contamination [14,15].

In addition, information from metabolomics provides additional insight into chemical contaminants in water that may influence bacterial diversity and would be useful to indicate the health status of an aquatic ecosystem and understand interactions between microbial communities and their environment [16,17]. For instance, a study in the Brisbane river (Australia) found that human interference was associated with a population size increment of Actinomycetes and Pseudomonadales, This increment was linked to higher levels of sugar alcohols, short-chain fatty acids and aromatic amino acids which contribute towards biofilm production [16].

In our work, we attempt to provide for the first time an overview of the microbiota from the Rimac river, comparing diversity patterns from Andean to Metropolitan areas and focusing on potential pathogenic bacteria.

## 2. Material and Methods

### 2.1 Area of study and collection

In January 2020, we sampled river water in 13 points along the extension of the Rimac river, within provinces of Huarochiri, Lima and Callao (Figure 1). Five sampling sites were from the Upper Rimac (1-5: Chicla, San Mateo City, Tamboraque, Huanchor and Chacahuaro); three, from the Middle Rimac (6-8: Santa Eulalia, Chaclacayo and Huachipa Bridge; and five, from the Lower Rimac near main bridges in the Metropolitan area (9-13: Libertadores, Nuevo, Universitaria, Faucett and Gambetta). Sterile plastic bottles were used to sample approximately 1.5 L of superficial water. When we sampled from a bridge, we attached the plastic bottle to another flask linked to a 30 m rope. In this case, we sampled superficial water (ca. 50 cm below the surface) from the middle of the bridge. Water samples were transported to the lab in coolers and were processed in the same day. We filtered river water using 0.2 µM Sterivex filters.

**Figure 1.**
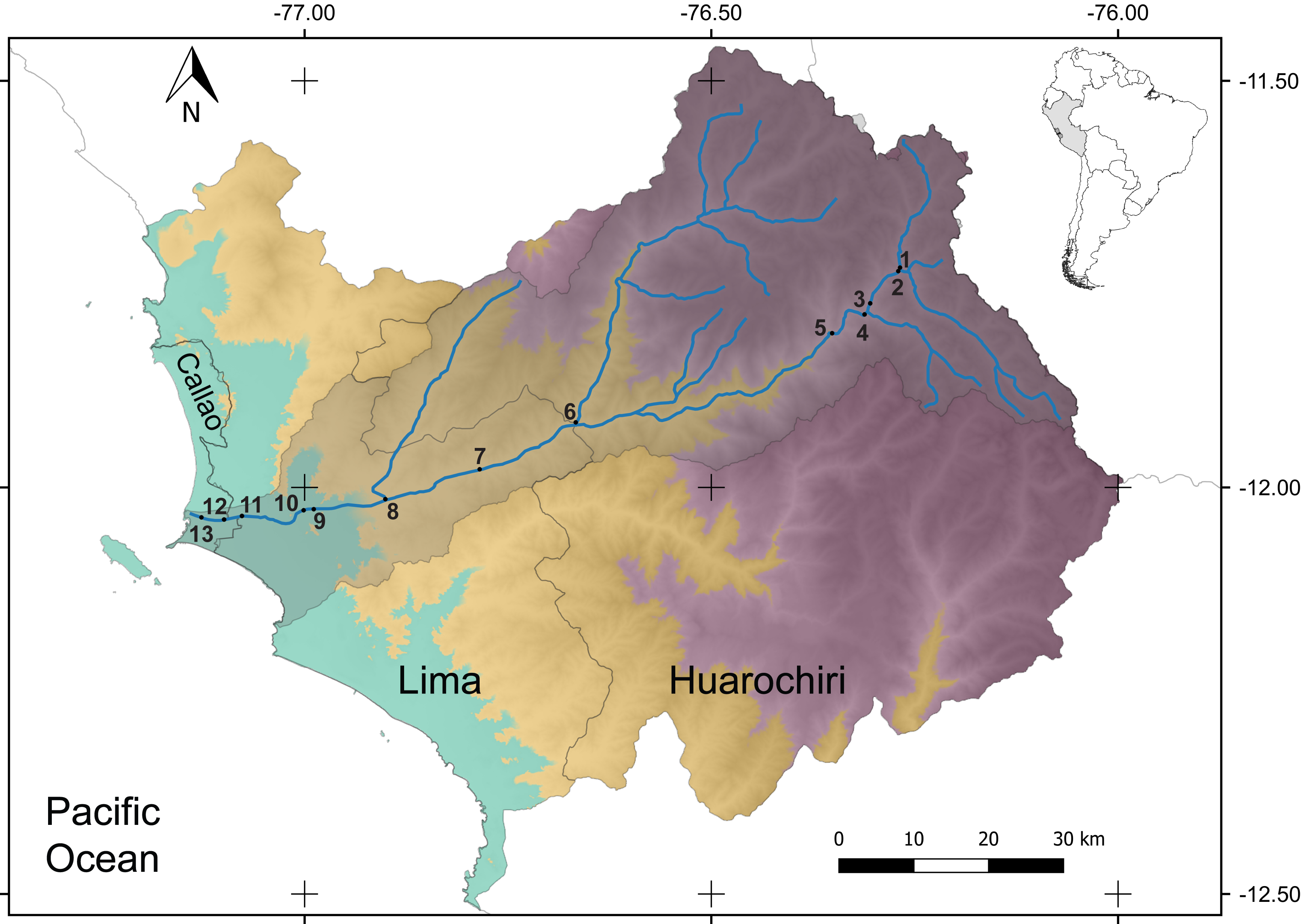
Area of study and sampling localities. The Rimac basin (in gray) is situated between the central Peruvian Andes and the Pacific Ocean. The river runs across Callao, Huarochiri and Lima provinces in the Lima region. Upper Rimac (>1 500 msl): 1, Chicla; 2, San Mateo; 3, Tamboraque; 4, Huanchor; 5, Chacahuaro. Middle Rimac (300 - 1 500 msl): 6, Santa Eulalia; 7, Chaclacayo; 8, Huachipa. Lower Rimac (Metropolitan area, 0 – 300 msl): 9, Libertadores; 10, Nuevo; 11, Universitaria; 12, Faucett; 13, Gambetta.

### 2.2 DNA isolation, amplification, and preparation of genomic libraries

Total DNA was isolated from the Sterivex filter using the PowerWater DNA kit (Qiagen) following manufacturer’s instructions. Then, we quantified DNA quality using a Qubit 3.0 (Thermo Fisher Scientific, USA) fluorimeter. For the preparation of genomic libraries, we followed the Illumina protocol [18] which consists in the amplification of the 16S rRNA V3-V4 region (ca. 460 nt). A second amplification is performed to attach oligo adapters (indexes) to each amplicon sample. Indexes will be informative to differentiate samples after the sequencing step. Each step was followed by amplicon cleaning using AMPure XP beads (Beckman Coulter, USA). In the final step, all samples were pooled and sequenced in an MiSeq (Illumina, USA) instrument. Raw sequences from the genomic libraries are deposited in the NCBI Bioproject database (accession number: PRJNA646070).

### 2.3 Bioinformatics and data analyses

Sequencing reads per each sample were analyzed using DADA2 [19] in R v3.6.3 [20]. We filtered reads using the parameter maxEE=2 for quality threshold as suggested before [21,22]. DADA2 was used to infer Amplicon Sequence Variants (ASV) which are groups of identical sequences. We also used DADA2 for taxonomic assignment, comparing ASV sequences against the ribosomal database SILVA v.138 [23] with the function “assignTaxonomy”. Identification up to the species (or multispecies) was done using the function “assignSpecies”. Then, we estimated Chao1, Shannon and Simpson alpha diversity indexes using the package phyloseq [24]. Significant differences among regions were evaluated using a paired Wilcoxon test. To estimate beta diversity, we normalized the ASV matrix using the variance stabilizing transformation (VST) method as suggested before for microbiome analyses [25]. This method is available in the package DESeq2 [26]. Then, we used a multidimensional scaling (MDS) approach to compare beta diversity among localities.

In addition, we used the taxonomic and occurrence information to produce a stack barplot of the most predominant classes. Then, we explored the most predominant genera using the package ampvis2 [27]. Next, we selected ASV with more than 1 000 counts and a taxonomic assignment until species level. We used this information to look for potential human and other animal pathogens in the Risk Group database of the American Biological Safety Association, specifically if species are pathogens for humans or other animals [28]. Finally, we used Piphillin [29,30] to predict functional content based on the frequency of the 16S rRNA sequences comparing them to annotated genomes in the Kyoto Encyclopedia of Genes and Genomes (KEGG) database. Predicted KEGG orthologues (KO) occurrence was retrieved from KEGG (October 2018) using a 97% cutoff threshold to create a gene feature table. We used the 20 most frequent unique KO per locality, then we grouped localities in three regions (Upper, Middle and Lower Rimac) and created a Venn diagram to look for shared KOs for each region using Venny [31].

### 2.4 Metabolomic analyses

Ten milliliters of each water sample and Milli Q water (Blank) were filtered through a Nalgene™ 0.22 µm syringe filter (Thermo Scientific) to remove suspended particulates. Each water sample was immediately extracted with dichloromethane (ACS grade, J.T. Baker) (3 times x 10 mL). After that, dichloromethane extracts were concentrated *in vacuo* and placed in a vial with 150 µL of dichloromethane and 2 µL of 26.7 ppb of toluene (GC grade, Sigma-Aldrich) (internal standard). The standard solution consisted in 2 ppm of toluene in dichloromethane, 2 µL of this solution was diluted with 150 µL dichloromethane. All water samples were analyzed except Santa Eulalia (6), Chaclacayo (7), Huachipa (8) and Gambetta (13) because these water samples were used completely in the step 2.1. Water samples, blanks and internal standards were analyzed by gas chromatography (GC) coupled to an APPI-Q-Exactive HF mass spectrometer (Thermo Fisher Scientific, USA). GC was equipped with a DB-5 column (30 m x 0.25 mm i.d., 0.25 μm thickness film). The oven temperature was programmed as follows: aliquots of 2 µL sample were injected at 45 °C for 2 min, increase at 10 °C/min until 270 °C, and hold for 7.5 min. Injections were made in splitless-mode with helium as the carrier gas (1.5 mL/min), injector temperature at 270 °C, and detector temperature at 270 °C. High-accuracy MS data were acquired in positive data-dependent acquisition (DDA) with scan range m/z 50– 750. Raw data from this GC-MS experiment was first converted into ABF format. Then, peak spotting was performed by exploring retention time and accurate mass. MS-DIAL v4.16 provided peak alignments of all samples and normalizes data based on TIC (total ion current). Final filtering was performed with a principal component analysis (PCA, p-anova < 0.05) using MATLAB vR2019b. Additionally, the software Compound Discovery was used for tentative identification using MS/MS data and comparing our results against several databases such as ChemBioFinder, Chemspider, Kegg, LipidBank, LipidMaps, Metlin and NIST. Besides, a PCA was performed to observe differences in mass profiles among samples group.

## 3. Results

### 3.1 Characterization of bacterial communities

16S rRNA amplicon sequencing produced an average of 265 625 [235 989-287 437] reads per sample. After discarding low quality sequences and merging forward and reverse reads, we obtained an average of 99 770 [76 129-122 999] final reads per sample (Supplementary Table 1). From this subset, we identified a total of 15 059 ASV. The first 151 ASV were frequent in more than 1 000 reads and the first eight, in more than 10 000 reads (Supplementary Table 2). Chao1, Shannon and Simpson diversity indexes were consistently higher in the Upper and Middle Rimac compared to the Lower Rimac (Figure 2). Alpha diversity values for each sample can be found in Supplementary Table 3, The information from the significance of pairwise comparisons can be found in Supplementary Table 4. In the MDS analysis, samples were grouped by their geographical location (Upper, Middle and Lower Rimac) except for Chacahuaro, a locality from the Upper Rimac that appeared closer to the Middle Rimac samples (Figure 3).

**Figure 2.**
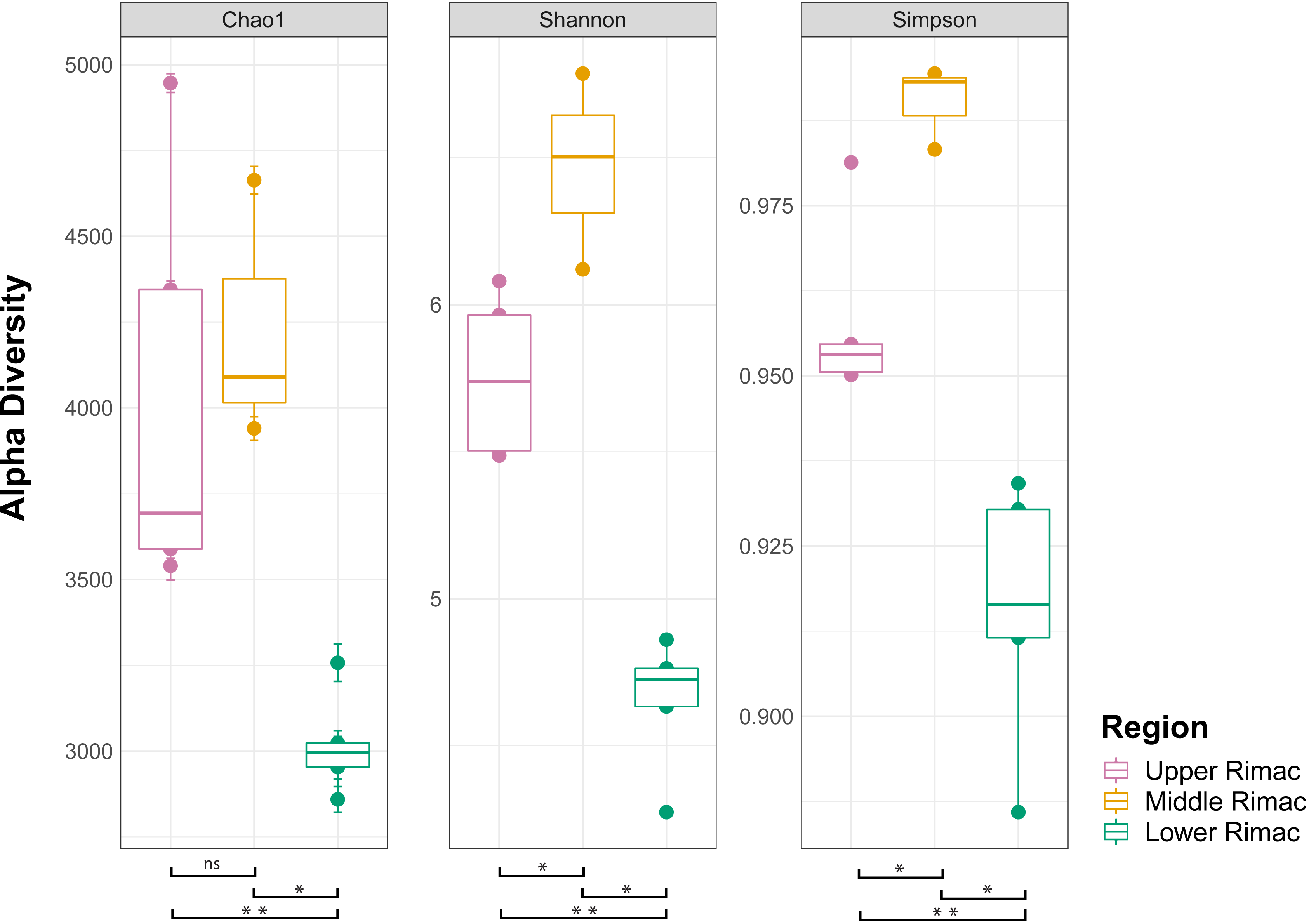
Alpha diversity indexes. Significant differences among the groups are denoted by ** (p<0.05), * (p<0.01), ns: non-significant.

**Figure 3.**
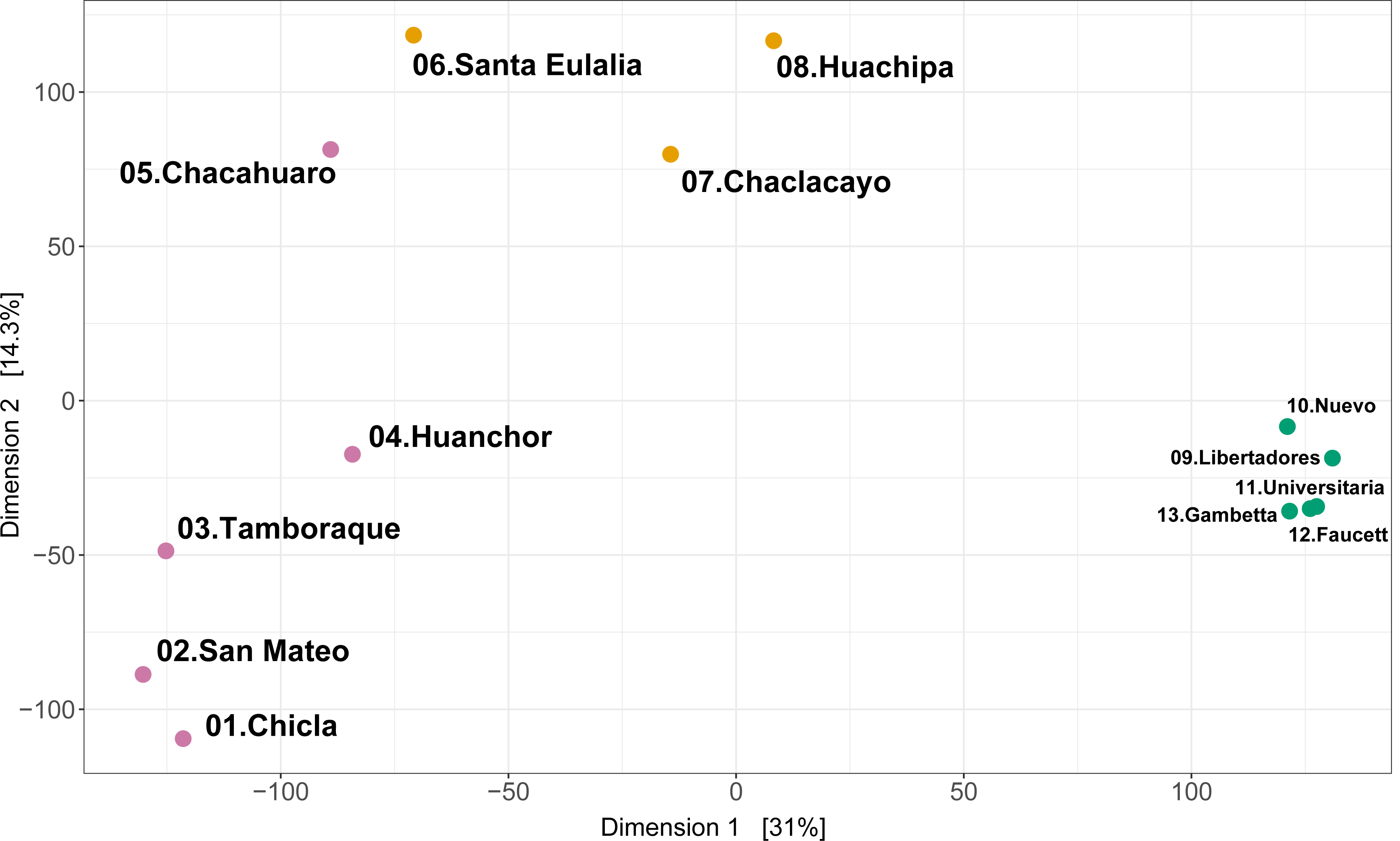
Multi-dimensional scaling (MDS). Principal coordinate analysis (PCoA) of ASV shows differences among groups of samples. Upper Rimac (light purple), Middle Rimac (orange), Lower Rimac (green).

Phyla Bacteroidota, Campilobacterota, Firmicutes, Fusobacteriota and Proteobacteria represented 74.79% (11 264 from 15 059) of the bacterial diversity in the ASV (Supplementary Table 5). Taxonomic identification of ASV was high for phylum, class, order, and family ranks (94 - 99%), and moderate for genus and species ranks (88 and 69%, respectively) (Supplementary Table 5). Classes Alphaproteobacteria, Bacteroidia, Campylobacteria, Clostridia, Fusobacteriia, and Gammaproteobacteria were the consistently common along all zones (Figure 4). Fusobacteriia declines in sampling sites 5 to 8 (Chacahuaro in the Upper Rimac zone and Middle Rimac samples). Campylobacteria especially rises in the Lower Rimac zone (Metropolitan area: sampling sites 9-13) but is also common in the Upper and Middle Rimac. The most frequent genera (Figure 5) in the Upper Rimac (sampling sites 1-4) were *Hypnocyclicus* (ASV2) and *Flavobacterium* (Bacteroidia), the latter was also predominant in the Middle Rimac. In the Lower Rimac, we found a dominance of *Arcobacter* (Campylobacteria). Human pathogenic bacteria identified until species level rank were *Arcobacter cryaerophylus* and *Prevotella copri* (Supplementary Table 6), and until the genus level rank were *Aeromonas, Escherichia, Pseudomonas* and *Shigella*. Other animal pathogens identified were *Flavobacterium succinicans* y *Faecalibacterium prausnitzii*.

**Figure 4.**
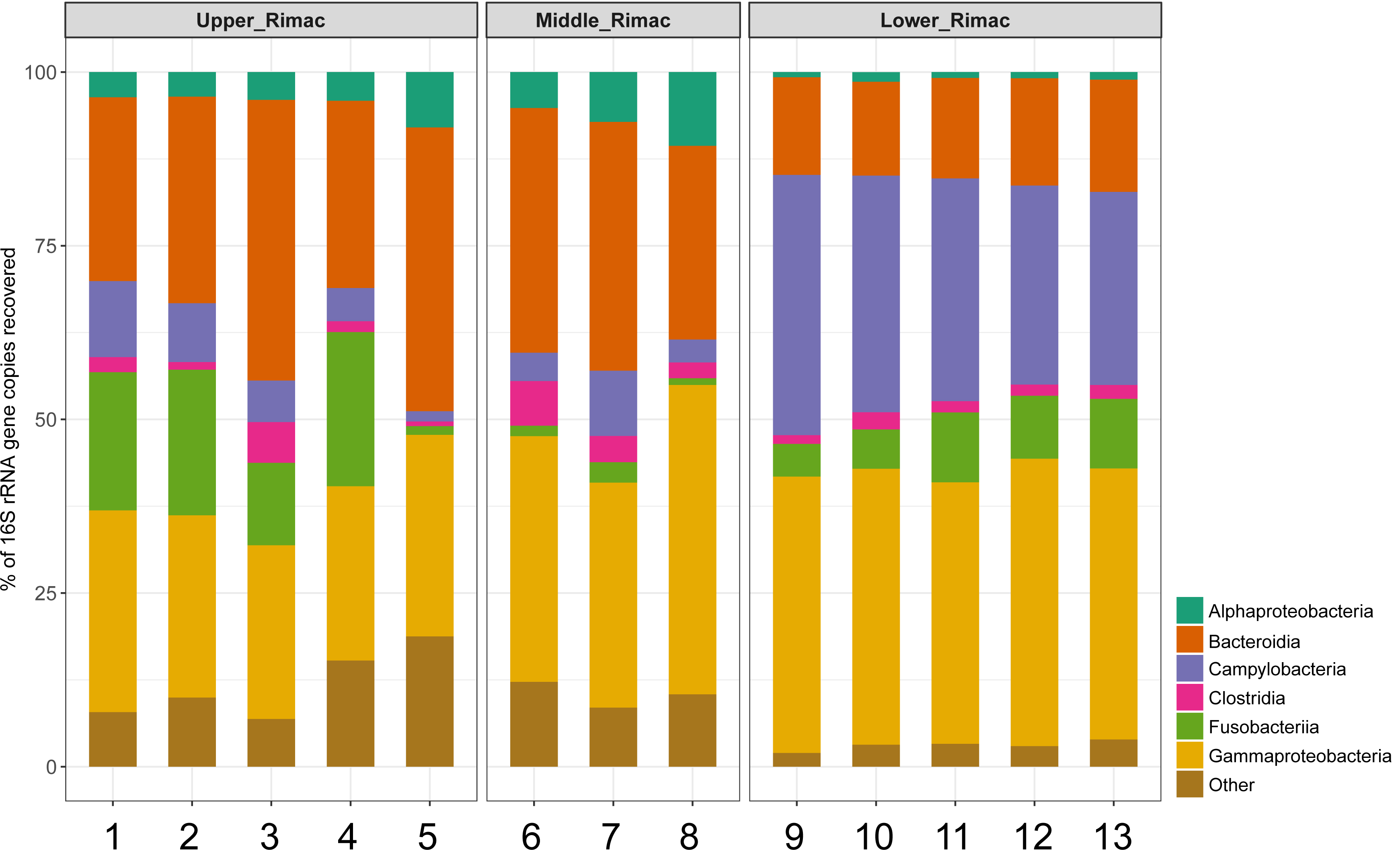
Stack barplot of the frequencies from the most common bacterial classes in each locality. Samples are numbered according to the information in Figure 1.

**Figure 5.**
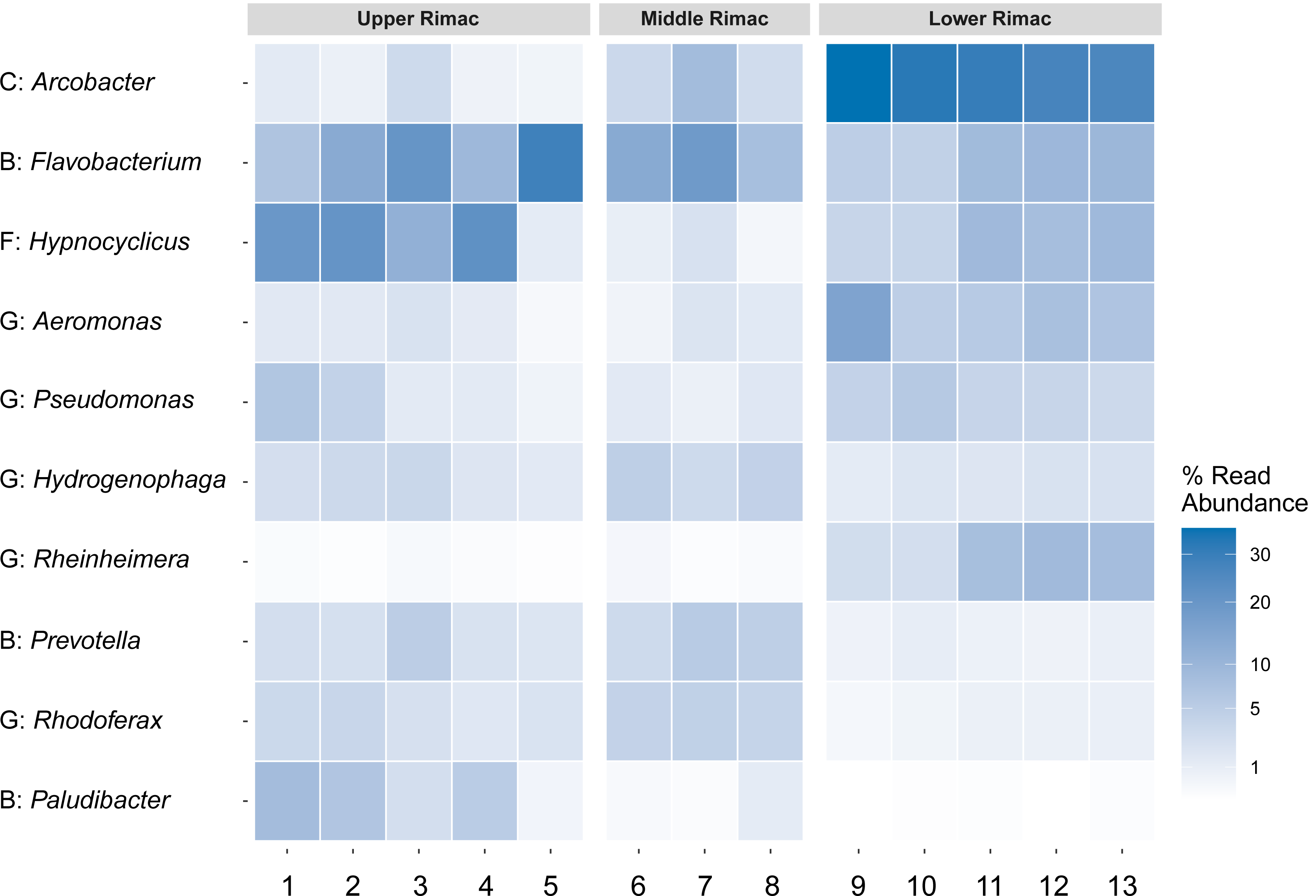
Heatmap of the abundance from the ten most frequent bacterial genera ordered from the most frequent to the less frequent. Classes codes: B, Bacterioidota; F, Fusobacteriia; G, Gammaproteobacteria. Samples are numbered according to the information in Figure 1.

Piphillin functional prediction of the top KEGG orthologies (KO) for each locality revealed 32 unique KO in the Upper Rimac, 26 in the Middle Rimac, and 27 in the Lower Rimac (Supplementary Table 7). The Venn diagram shows nine KO shared by the three zones, 15 KO were only shared between the Upper and Middle Rimac and four, only between the Upper and Lower Rimac (Supplementary Figure 1). The most abundant predicted pathways shared by the three zones (Supplementary Table 8) were related to bacterial chemotaxis (methyl-accepting chemotaxis protein/MCP, K03406), metabolism of xenobiotics (glutathione S-transferase, K00799), and aminoacyl-tRNA biosynthesis (K14218, K14228). Between the Upper and Middle Rimac the most frequent pathways predicted were ATP-binding cassette transporters and quorum sensing (Leucine-Isoleucine-Valine *liv* gene cluster, K01996 - K01999).

### 3.2 Metabolomic analyses

Results from the GC-MS experiment can be found in Supplementary Table 9. PCA performed from these results (Supplementary Figure 2), first showed that the Upper and Lower Rimac were different from blank and internal standard samples. Some localities from the Upper zone (Tamboraque, Huanchor and Chacahuaro) were well differentiated from the other areas. However, other Upper Rimac samples (Chicla and San Mateo) were clustered within the Lower Rimac because of similar chemical composition. Additionally, GC-MS (APPI) in positive mode allowed to detect eleven compounds (p-anova < 0.05, total score > 95%) using Compound Discovery and MS-DIAL softwares (Supplementary Table 10). The loading plot of PCA coefficients from metabolomics analysis (Supplementary Figure 3) showed that the Upper Rimac localities, Tamboraque, Huanchor and Chacahuaro are influenced by the following compounds: nanaomycin derivates, natural bacterial antibiotics, and hispidulin, a natural plant flavonoid (Supplementary Table 10). Whereas the other Lower Rimac samples were influenced by propazine-2-hydroxy, an herbicide. Additionally, a natural flavonoid, isopeonol, and other pharmaceutical compounds such as desloratadine, meprylcaine and N-(4-ethoxyphenyl)acetamide were identified in both zones (Supplementary Table 10).

## 4. Discussion

Our study provides the first overview of the bacterial community diversity in the Rimac river (Lima, Peru) using a 16S rRNA amplicon sequencing approach. We described bacterial community shifts along an altitude gradient from the Upper Rimac zone (1 500 – 4 500 msl) to the Middle (300 – 1 500 msl) and Lower Rimac zones (0 – 300 msl). The Lower Rimac crosses the Lima Metropolitan area, a mega-city situated on the hyper-arid Peruvian desert [4,32], being an essential water source for this city.

Previous studies in rivers have shown different patterns of diversity along altitudinal gradients. For instance, a diversity decline downriver was found in the Danube in free-living and particle-associated communities [12], while an increase was found in the Mississippi especially in organisms related to suspended particles [33]. In our case, different estimators of alpha diversity (Chao1, Shannon and Simpson) provided evidence of a decline of diversity in the Lower Rimac (Metropolitan area). The greatest mean diversity values were found in the Middle Rimac and may be influenced by confluences with minor rivers such as the Santa Eulalia river (sampling site 6). It appears necessary to sample more localities between sites 5 to 8 to provide a better assessment of local diversity in the Middle Rimac. Besides, we sampled in January 2020, rainy season in the Andes, so it would be desirable also collecting during the dry season to compare temporal diversity patterns.

Bacterial communities in high-altitude aquatic environments are still not thoroughly studied. We found only few characterizations using 16S rRNA sequencing, especially in lakes. For instance, community profiling of Tibetan and Pyrenean lakes showed dominance of classes Actinobacteria, Alphaproteobacteria, Betaproteobacteria, and phyla Bacteroidota [34–36]. Our samples did have a major presence of the Bacteroidia class, part of the latter phylum, and class Alphaproteobacteria. However, we found no Betaproteobacteria. Additional surveys in high-altitude rivers are necessary to have a broader view of bacterial communities in these kinds of habitats.

On the other hand, samples from the Lower Rimac looked more similar among each other than samples in the Upper or Middle Rimac (MDS, Figure 3). Community composition in the Lower Rimac is probably influenced by similar pollution constrains such as human feces and domestic sewage that are produced in the Metropolitan area [37].

Furthermore, we found a constant occurrence of the phyla Bacteroidota, Campilobacterota, Fusobacteriota and Proteobacteria along the Rimac river basin. This result is similar to a previous report in the polluted Pinheiros river in Sao Paulo, Brazil [14]. The authors associated the high presence of these phyla with freshwater environments and domestic sewage sludges. In addition, they also found *Arcobacter cryaerophilus*, as the predominant species in the river. This species has been reported as very frequent in other rivers such as the Yangtze (China) [37], many rivers in Nepal [38], and the Llobregat (Spain) [39]. As mentioned before, *Arcobacter* is an emergent enteropathogen and indicator of fecal contamination. It can also occur in water treatment and sewage systems [40] and survive adverse conditions imposed by food processing and storage [41]. *Arcobacter* is also abundant in effluent from wastewater treatment plants [42]. In Lima, the main water treatment plant is situated in the Lower Rimac nearby the sampling site 9 (Figure 1). Thus, our results corroborate previous reports of the endurance of this genus and the necessity of a better water quality assessment based in amplicon sequencing [42] not only for environmental samples but also for water samples taken after treatment.

Moreover, *Arcobacter* has shown resistance to several antibiotics [41] and it has been proposed to be involved in the exchange of resistance genes between gram-negative and gram-positive phyla [43]. Notwithstanding, reports in Peru on the pathogenicity of this genus are scarce, e.g. we found only one published study that mentioned *Arcobacter* in stool from children with diarrhea [44]. In particular, *Arcobacter cryaerophilus*, the most frequent ASV in the Metropolitan area, has been associated with severe diarrhea and found to carry several virulence genes [45].

In the Upper Rimac, we found a predominance of *Hypnocyclicus* and *Flavobacterium*, the latter has been related to several issues in fish such as “cold water” and *columnaris* disease [46]. In particular, *Flavobacterium succinicans*, the most frequent *Flavobacterium* (Supplementary Table 5), has been associated to bacterial gill disease in trout [47].

The high occurrence of bacterial chemotaxis pathway, a necessary function to move towards nutrients or away toxins, would be connected to bacterial growth and survival in aquatic environments [48]. A study in the Pinheiros river found similar high occurrence of bacterial chemotaxis and flagellar assembly functions and hypothesized that this could be influenced by the elevated concentration of nutrients, e.g. ammonia and phosphates [14]. In our study, the most predicted KO in our samples was the MCP protein (Supplementary Table 8). This protein is important for bacterial motility because it triggers the activation of flagella, and has been also encountered in other highly polluted rivers such as the Yamuna, a major tributary of the Ganges river in India [49]. In addition, we also found a high occurrence of predicted ABC transporters which are ubiquitous proteins involved in several processes including nutrient uptake and chemotaxis [50]. According to the KEGG annotation these proteins also take part in quorum sensing functions. Quorum sensing has been profusely studied in free-living bacteria in laboratory conditions. However, it is now known that, in nature, bacteria can use quorum sensing mechanisms in fluid environments such as rivers, streams, intertidal and marine areas by forming biofilms [51]. Another abundant function was related to the Glutathione metabolism. Glutathione S-transferase is involved in biodegradation of xenobiotics, defense against chemical and oxidative stress, and antibiotic resistance [52]. Rivers have been shown as reservoirs of genes related to antibiotic resistance influenced by anthropogenic causes [49,53]. Further research is necessary to obtain environmental metagenomes from the Rimac and to look for possible genes linked to resistance to antibiotics in the Lower and Middle Rimac or resistance to heavy metals which may be frequent in the Upper Rimac.

Metabolomics analysis by GC-MS showed differences in chemical composition between some localities from the Upper Rimac with respect to the Lower Rimac. This variance may be due to less population density in the Upper Rimac and distance from the Metropolitan area. However, we also found that some localities from the Upper Rimac are as contaminated as the Lower Rimac. This apparently contradictory results highlight that more work should be done to understand the pollution patterns caused by human influence. For instance, we found signals of some pain-relieving compounds in the Upper and Lower Rimac but no signal of synthetic antibiotics. Nanaomycin derivates are actinomycete metabolites, biosynthesized by *Streptomyces* [54]. These compounds had a higher frequency in the Upper Rimac. However, we did not identify any *Streptomyces* in our data (Supplementary Table 5). This could be because *Streptomyces* are generally found in the river sediment [55,56]. Further sampling efforts should also consider different portions of the water column and sediment.

Finally, our study could be complemented with metabarcoding analyses of other pathogenic protozoa [7] or invertebrates which are also used as indicators of water quality [57]. We believe that this initial work will promote more similar studies in our country. Further work that integrates -omics and environmental sciences with public health studies will be beneficial for future public policies especially focused on rural and sub-urban areas in Lima which depend on the Rimac river water.

## Ethical approval

The study did not involve experiments in humans nor animal subjects.

## 5. Acknowledgments

PER thanks to Yulissa Estrada and Grecia Valdivia (Universidad de Ingeniería y Tecnología. Lima, Peru) for their support during sampling in the Upper Rimac. In addition, PER thanks to Raul Condori and Enrique Santisteban for their kind support during sampling in the Metropolitan area. AJI thanks Thermo Fisher Scientific for its support in the metabolomic analyses. PER and PW thank to Arturo Gonzáles and Stefany Infante (Universidad de Piura) for their help in the lab work. All authors thank to Guillermo Trujillo (GenLab) for his support in the preparation of genomic libraries.

Supplementary material can be requested to http://pedro.romero@upch.pe

